## Supplementary file for "ATLASx: a computational map for the exploration of biochemical space"

### Supplementary Tables

**Supplementary Table S1:** Import and curation of compounds from different sources.

| | | Database | Description | Collected | Imported | Unique in source database | Total unique compounds |
|---|---|---|---|---|---|---|---|
| Compound DBs | biological | MetaCyc | Manual/Cpds of sequenced organisms | 15,819 | 14,828 | 12,524 | |
| | | Model SEED | Manual/KEGG and GSMs | 33,995 | 20,665 | 17,132 | |
| | | KEGG Comp. | Manual/Cpds & biopolymers relevant to biology | 18,625 | 17,397 | 15,064 | |
| | bioactive | KEGG Drug | Manual/approved drugs in Japan, USA, & Europe | 11,140 | 7,766 | 4,514 | |
| | | Drugbank* | Approved drugs + discovery-phase drugs | 8,350 | 6,279 | 3,850 | |
| | | ChEBI | Chemical Entities of Biological Interest | 56,530 | 32,691 | 29,080 | |
| | | HMDB | Small cpds found in the human body | 228,017 | 177,096 | 98,400 | |
| | | MetaNetX** | The metabolites in the GSMs + other databases | 200,132 | 183,788 | 87,464 | |
| | | ChEMBL | Manual /bioactive/drug-like cpds | 1,727,112 | 1,595,615 | 1,365,379 | |
| | | **Total** | | **2,297,709** | **2,056,125** | **1,633,407** | **1,500,222** |

* Experimental drug    ** Lipids excluded    Cpd: Compound

**Supplementary Table S2:** Import and curation of reactions from different sources.

| | Database | Description | Collected | Imported | Unique in source database | Total unique reactions |
|---|---|---|---|---|---|---|
| Reaction databases | HMR | GSMs for human metabolic reactions | 8,182 | 5,108 | 4,257 | |
| | MetaCyc | Manual/reactions in pathways of sequenced orgs | 16,052 | 15,438 | 12,093 | |
| | KEGG | Manual/reactions in KEGG enzyme or KEGG pathway | 10,829 | 10,685 | 10,338 | |
| | MetaNetX | The reactions in the GSMs + other databases | 42,182 | 40,767 | 25,871 | |
| | Reactome | Manual/reactions in human | 1,872 | 1,568 | 777 | |
| | Rhea | Manual curation of biochemical rxns/cpds from ChEBI | 20,770 | 19,325 | 11,753 | |
| | Model SEED | Manual/KEGG and GSMs | 44,031 | 44,010 | 25,807 | |
| | BKMS | Rxns of BRENDA, KEGG, MetaCyc, and SABIO-RK | 31,740 | 18,139 | 17,409 | |
| | BiGG models | Manual/reactions from GSMs | 28,299 | 16,581 | 8,354 | |
| | Brenda | Large set of enzyme functional data | 31,741 | 9,214 | 6,578 | |
| | **Total** | | **235,698** | **180,835** | **123,237** | **56,087** |

GSM: Genome scale models Rxn: Reaction Cpd: Compound

**Supplementary Table S3:** Quality of reactions in different sources based on mass balance and EC annotation.

| Database | # Total unique reactions | # EC annotated reactions | # Balanced* reactions | # Balanced & EC annotated reactions |
|---|---|---|---|---|
| KEGG | 10,338 | 9,996 | 7,789 | 6,859 |
| Brenda | 6,578 | 5,982 | 5,855 | 5,281 |
| Rhea | 11,753 | 8,711 | 8,972 | 6,406 |
| BiGG | 8,354 | 3,657 | 3,972 | 1,755 |
| Model SEED | 25,807 | 8,662 | 16,041 | 6,300 |
| MetaNetX | 25,871 | 13,288 | 15,589 | 8,441 |
| MetaCyc | 12,093 | 8,495 | 8,458 | 6,282 |
| HMR | 4,257 | 3,153 | 2,969 | 2,001 |
| Reactome | 777 | 333 | 466 | 200 |
| BKMS | 17,409 | 14,589 | 10,962 | 9,252 |

* removed isomerases, transports

**Supplementary Table S4:** Network statistics of bioDB, bioATLAS, and chemATLAS networks.

| Network | Property | bioDB | bioATLAS | chemATLAS |
|---|---|---|---|---|
| Weighted network | Number of nodes | 14,914 | 844,337 | 1,876,992 |
| | Number of edges (CAR > 0) | 62,299 | 2,503,627 | 5,717,409 |
| Unweighted network (Only edges with CAR > 0.34) | Number of nodes | 14,084 | 617,942 | 1,854,423 |
| | Number of edges (CAR > 0.34) | 25,624 | 982,343 | 2,778,445 |
| | Number of components (disjoint graphs) | 623 | 68,912 | 151,390 |
| Biggest component | Number of nodes | 12,434 | 361,405 | 1,264,423 |
| | Number of edges | 24,575 | 774,584 | 2,297,335 |
| | Percent of total number of nodes | 88.28% | 58.49 % | 68.19 % |
| | Percent of total number of edges | 95.84 % | 78.85 % | 82.68 % |
| | Diameter [a] | 32 | 27 | 46 |
| | Average path length [b] | 7 | 9 | 12 |

[a] Length of longest shortest path between any two nodes, [b] Length of average shortest path length between any two nodes

**Supplementary Table S5:** Reconstruction of known bioDB reactions within ATLASx

| | | Number of reactions |
|---|---|---|
| **Total reactions bioDB** | | 56,087 |
| **Filtered bioDB reactions** (only reactions with defined molecular structures of reactants are kept) | | 41,680 |
| **Reaction reconstruction** | **Exact coverage:** Reactions reconstructed with BNICE.ch rule | 11,172 |
| | **Alternative cofactor usage:** 1-step reconstruction of main biotransformation(s) within ATLASx | 14,193 |
| | **2-step reconstruction:** Main biotransformations reconstructed in max. of 2 reaction steps | 2,625 |
| | **3-step reconstruction:** Main biotransformations reconstructed in max. of 3 reaction steps | 1,175 |
| | **4-step reconstruction:** Main biotransformations reconstructed in max. of 4 reaction steps | 603 |
| **Total number of reconstructed bioDB reactions** | | 29,768 |
| **Percentage of reconstruction in filtered bioDB reactions** | | **71.42** |

### Supplementary Figures

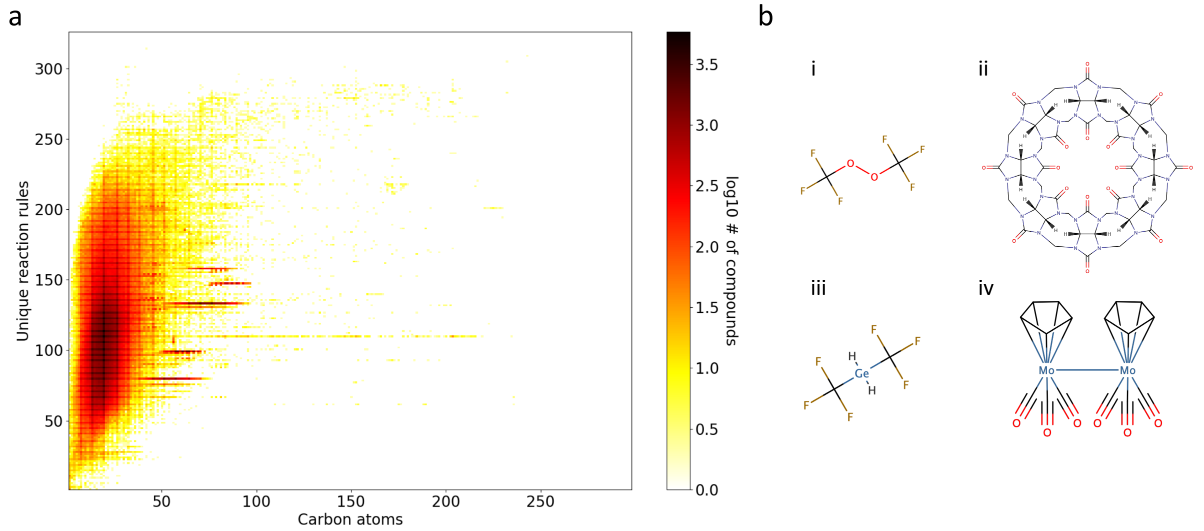


**Supplementary Figure S1: Reactive site analysis of bioATLAS compounds. a**, Heatmap showing the distribution of compounds as a function of their number of carbon atoms versus the number of reaction rules assigned to them. Darker colors indicate a higher number of compounds on a logarithmic scale. **b**, Examples of four bioactive compounds for which BNICE.ch could not find any reactive site. i, Bis(trifluoromethyl)peroxide(BTP), ii, cucurbit[8]uril, iii, Bis(trifluoromethyl)germane, iv, bis[tricarbonyl(η5-cyclopentadienyl)molybdenum](Mo—Mo).

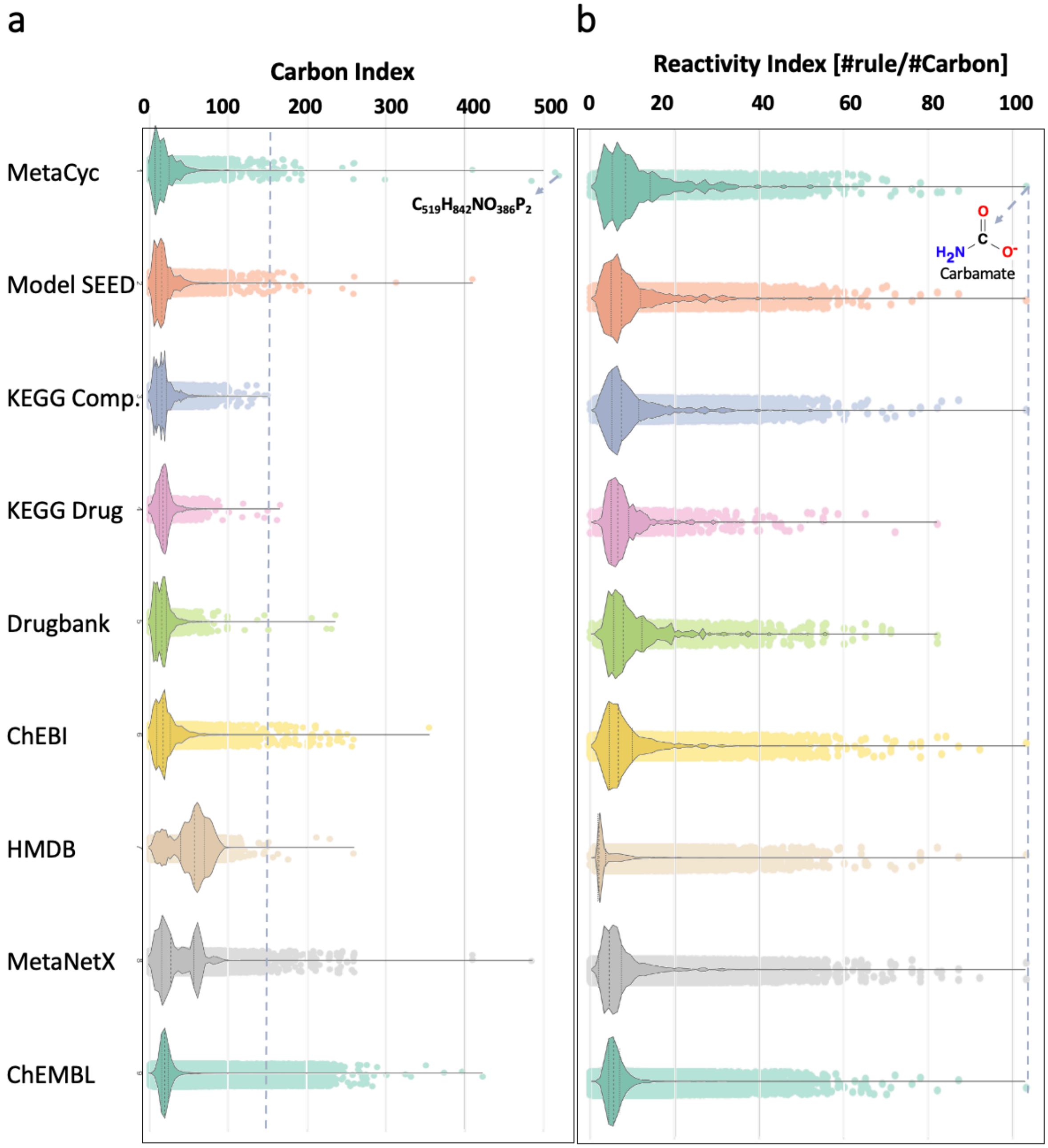


**Supplementary Figure S2: Biochemical reactivity of compounds compared across different databases.** For each database, the distribution of compounds is represented as a violin plot. **a,** The carbon index (i.e., number of carbon atoms inside the molecule) ranges from 0 to 511. The dashed line shows the maximum of carbon index in compounds of KEGG database, indicating broader distribution of molecules among bioactive molecules. **b,** The reactivity index is calculated as the number of reaction rules assigned to a given compound, divided by the number of carbon atoms within the molecule. The reactivity index ranges from 0 to 104.

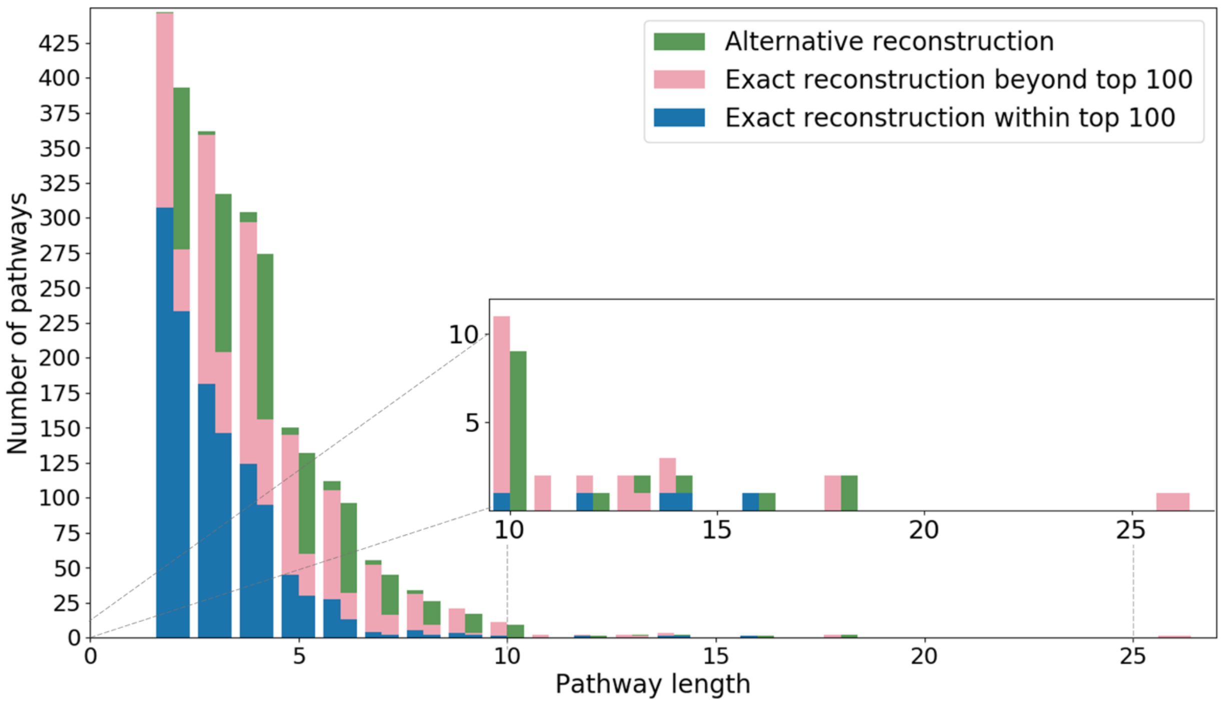


**Supplementary Figure S3:** Distribution of the length (i.e., number of reaction steps) of reconstructed MetaCyc pathways. Only bioDB pathways with complete reaction mechanisms are shown on the right side of the bar and all bioDB pathways are shown in the left side of the bar.

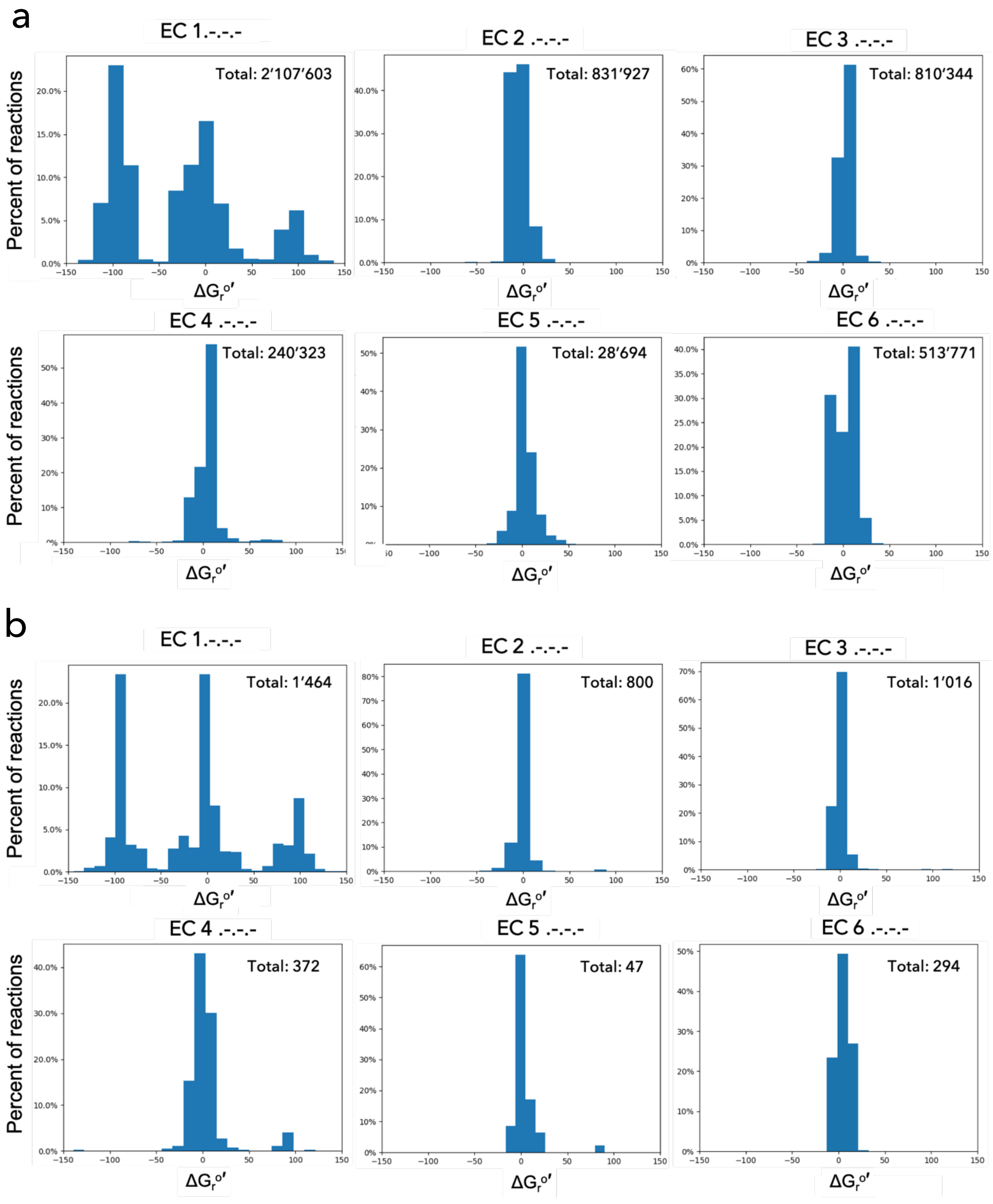


**Supplementary Figure S4:** Histogram of the distribution of standard Gibbs energy of reaction for each 1st level EC class. In the upper right corner total number of reactions plotted indicated. a, distribution of the Gibbs free energies calculated within chemATLAS space (total 4’231’154 (81%) reactions of chemATLAS have Gibbs free energy). b, distribution of the Gibbs free energies calculated within bioDB space (total 3’809 reactions have Gibbs free energy estimation).

*Note that total number of the reactions that have a BNICE.ch reaction rule and energy estimation is not equal to the sum of totals per EC class as same reaction can have more than one first level EC class assigned. Besides this, not all reactions that have BNICE.ch rule assigned can have an energy estimation as energy cannot be estimated for reactions including generic compounds.
